# Nematicidal indole oxazoles and chemoattractants from soil bacteria

**DOI:** 10.64898/2026.01.20.700618

**Authors:** Kaetlyn T Ryan, Julia M Duncan, Chris S Thomas, Marc G Chevrette, Martel L DenHartog, Kristin J Labby, Joanna Klein, Mostafa Zamanian, Jo Handelsman

**Affiliations:** Department of Pathobiological Sciences, University of Wisconsin-Madison, Madison, WI, USA; Wisconsin Institute for Discovery, University of Wisconsin-Madison, Madison, WI, USA; Department of Chemistry and Biochemistry, University of Alaska Fairbanks, Fairbanks, AK, USA; Research and Development, Forensic Fluids Laboratory, Plymouth MN, USA; Department of Chemistry, Beloit College, Beloit, WI, USA; Department of Biology, University of St. Thomas, St. Paul, MN, USA; Department of Plant Pathology, University of Wisconsin-Madison, Madison, WI, USA

## Abstract

Ecological interactions between bacteria and nematodes in many environments provide a basis for the prediction that diverse bacteria produce anti-nematode compounds. The discovery of microbial secondary metabolites with broad-spectrum nematostatic or nematicidal properties can be hastened by drug screening approaches that include several nematode species and phenotypes. We cultured a collection of 22 soil-derived bacterial isolates that carry in their genomes putative pathways for production of unknown secondary metabolites. Isolates were cultured in various media to enhance natural product diversity and yield, and we evaluated culture filtrates for activity against two evolutionarily distinct nematode species: Clade V free-living nematode *Caenorhabditis elegans* and Clade III mammalian parasitic nematodes in the genus *Brugia*. Partitioned extracts from *Pseudomonas* sp. strain TE4607 stunted *C. elegans* development and caused motility defects in both blood-circulating larval and adult stages of *Brugia*. The primary active compound was identified as labradorin 1, an indole with known antibacterial and anticancer properties that had not been previously described as affecting nematodes. Notably, filtrates of *Pseudomonas* sp. TE4607 cultures attracted free-living nematodes in sensory assays, adding to evidence that certain *Pseudomonas* species modulate the behavior of free-living nematodes. These findings underscore the need to further explore the link between nematode sensory responses and whole-organism effects of microbial metabolites, with potential applications in anthelmintic discovery.

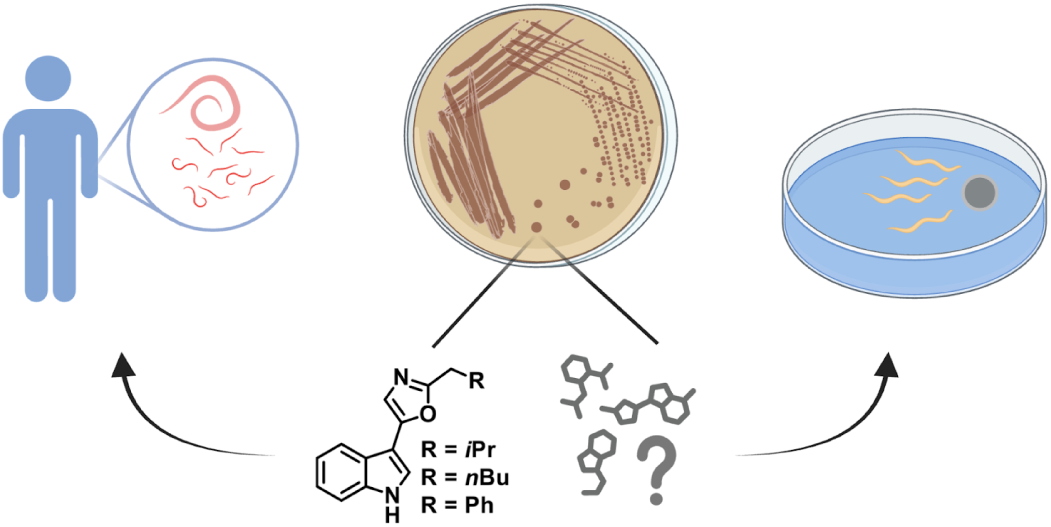

## Introduction

Bacteria and nematodes are ubiquitous across diverse environments and can engage in competitive or antagonistic interactions. Free-living nematodes participate in complex interactions with bacteria and fungi, including predator-prey dynamics^1^ and the exchange of toxins^2–4^. Nematodes possess a remarkable repertoire of chemosensory receptors^5^, and many microbes produce signals that influence nematode sensory behaviors^6^. Previously characterized microbial cues include metabolites that facilitate nematode food-seeking or pathogen avoidance^7–9^ and attractants used to entrap or infect nematodes^10–12^. These inter-kingdom dynamics have the potential to provide evolutionary pressure for microbes to develop metabolites or other molecular machinery with both nematicidal and sensory-modulating properties.

Bioactive compounds emerging from these microbe-nematode interactions hold promise as pharmaceutical treatments for parasitic diseases and as biocontrol agents targeting closely related free-living plant-parasitic nematodes. Ivermectin and emodepside are examples of antiparasitic drugs derived from microbial natural products^13,14^ that exhibit activity against nematode species spanning several clades, owing to significant genomic conservation within the nematode phylum.

Parasitic nematodes infect billions of people, cause significant suffering in companion animals, present zoonotic risks, and inflict major economic losses through their impacts on livestock and crops^15–17^. For example, lymphatic filariasis, caused by *Brugia malayi*, is a CDC-priority neglected tropical disease (NTD) and a target for global elimination by the WHO. The need for novel treatments and therapies to address nematode infections is urgent given existing and emerging resistance to antiparasitic drugs in current use^18–20^ and shifts in vector prevalence and agricultural conditions caused by climate change. Despite the promise of microbial secondary metabolites as a source of new nematicides or antiparasitics, efforts to strategically optimize and scale screening methods face several challenges. Free-living nematodes offer a tractable and scalable model in which to screen for bioactivity, but this activity does not reliably predict efficacy against parasitic nematodes^21–23^. Direct evaluation of anthelmintic potential against parasitic nematodes is preferred, but such efforts are complicated by the need to propagate the nematodes in vertebrate and invertebrate hosts^24^. Moreover, the possibility of rediscovering known antiparasitics adds significant investment risk to the search for nematicidal natural products.

Here, we attempt to address some of these challenges by performing parallelized screening of phylogenetically distant free-living nematodes (*Caenorhabditis elegans*, Clade V) and mammalian parasitic nematodes (*Brugia spp.*, Clade III). We focused on accessible parasite life cycle stages of *Brugia*, which can serve as predictive proxies for activity against medically relevant but lower-throughput stages in a two-tiered approach^23^. To enhance the chance of discovery of new compounds, we tested bacterial isolates that were prioritized based on genomic analysis for unusual biosynthetic capacities and grown in diverse culture media. We evaluated several phenotypic endpoints relating to nematode development, motility, tissue toxicity, and sensory responses. A deeper understanding of how nematodes detect and respond to microbial cues may reveal novel lead compounds and inform future strategies for nematicide and anthelmintic discovery.

## Results and Discussion

### Identification of bacterial isolates with nematicidal and anthelmintic activity

A large library of bacterial isolates was generated from soil samples collected from a variety of locations in Wisconsin, Illinois, and Minnesota. The bacterial isolates were screened by students for antibacterial activity and then included in the Tiny Earth collection^25^. Twenty-two isolates were prioritized based on antiSMASH sequence analysis indicating that they likely had unstudied biosynthetic gene clusters predicted to be responsible for synthesis of bioactive small molecules. Bacterial genera represented in this diverse collection included *Pseudomonas*, *Flavobacterium*, *Paraburkholderia*, *Curtobacterium*, *Streptomyces*, *Paenarthrobacter*, *Providencia*, *Bacillus*, *Arthrobacter*, and *Paenibacillus*. We performed primary screens of culture supernatant filtrates for each isolate grown in four media (M9, PDB, LB, and TSB10) to broaden the range of secreted natural products^26^.

These filtrates were first screened against model nematode *C. elegans* (Clade V) and parasitic *Brugia* microfilariae using three phenotypic endpoints: development, motility, and tissue toxicity^23^ (**Figure 1A**). In these high-content imaging assays, four technical replicates (i.e., populations in a microtiter plate well) were performed for each isolate, and phenotypes were quantified and normalized using image-processing software^27^. The measurements collected from these analyses include worm size as a proxy for *C. elegans* development, optical flow to quantify parasite motility, green fluorescence (RFU) as an indicator for tissue toxicity and reductions in parasite viability, and progeny quantity to describe fecundity. The positive control in the development assay was 50 µM albendazole sulfoxide, which restricts larval development to the L2 phase. The positive control for parasite assays was heat killing, which abolishes motility and generates the maximum achievable tissue toxicity fluorescence value for a given well. In this initial screen, filtrates that elicited phenotypes most similar to positive controls were selected for further analysis.

**Figure 1.**
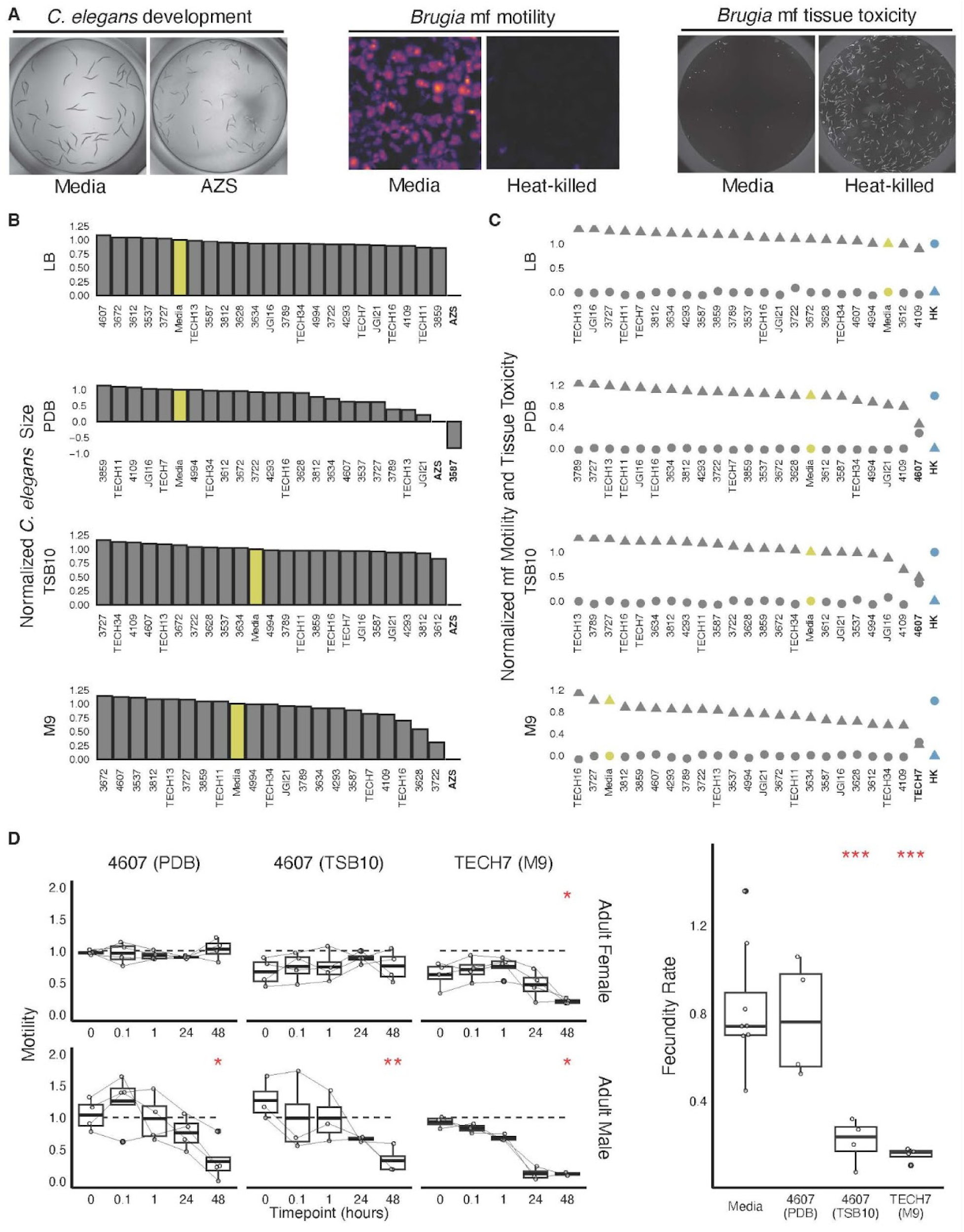
Primary nematode phenotypic screen of 22 bacterial isolates grown in four media conditions. **(A)** Representative images of controls 48 hours post treatment across primary phenotypic endpoints. Left: *C. elegans* development assay showing worm size after media (− control) and albendazole sulfoxide (AZS, + control) treatment. Middle: *Brugia* microfilariae motility assay showing optical flow heat maps of media treated (-control) and heat-killed (+ control) worms where color is brighter with increased motility. Right: *Brugia* microfilariae tissue toxicity assay showing staining with Promega’s CellTox Green reagent in media (− control) and heat-killed (+ control) worms where fluorescence indicates decreased viability. **(B)** Effects of test isolates on *C. elegans* development. Mean worm sizes of treated worms normalized between positive (blue: 50 µM AZS) and negative (yellow: media) controls. **(C)** Effects of test isolates on *Brugia* microfilariae (mf) motility (triangular points) and tissue toxicity (circular points). Mean phenotypic values for test strain wells are normalized between positive (blue: heat-killed) and negative (yellow: media) controls. **(D)** Effects of top microfilariae hits *Pseudomonas* sp. TE4607 and *Pseudomonas viciae* TECH7 on *Brugia* adult motility (left panel) and fecundity (right panel). Supernatants were made from cultures grown in media conditions that elicited the most potent effects in the primary microfilariae screen. Individual worm motility at each time point is normalized to mean control (media) values and fecundity at 48 hours is normalized to initial time point values for each worm. *C. elegans* and microfilariae phenotypes were normalized as follows: (X − positive control) / (negative control − positive control) where X is the phenotypic endpoint value while adult parasite data points were normalized to DMSO and media controls alone: X / negative control after normalizing to individual initial motility scores. Statistical analyses were performed via t-test and reported as follows, * : p<0.05, ** : p<0.01, *** : p<0.001, **** : p<0.0001.

Of the 88 filtrates tested, only those from *Flavobacterium* sp. TE3587 grown in PDB media inhibited the growth of *C. elegans* larvae at or beyond the level of positive controls (**Figure 1B**). The absence of other positive hits may be due to low concentrations of active compounds in the unconcentrated filtrate, the stringency of the required activity level, or the low permeability of the *C. elegans* cuticle^28,29^. The *Flavobacterium* sp. TE3587 genome contains several lanthipeptide biosynthetic gene clusters that could be responsible for this activity. However, this isolate was deprioritized due to lack of broad-spectrum activity, as these filtrates did not cause *Brugia* microfilariae motility defects or tissue toxicity (**Figure 1C**).

*Brugia* microfilariae screens revealed two additional isolates that reduced motility and caused tissue toxicity: *Pseudomonas* sp. TE4607 grown in PDB and TSB10 and *Pseudomonas viciae* TECH7 grown in M9. These results were replicated across two batches of parasites reared separately. The origin of each of these strains is described in methods. Next, we evaluated the effects of these active isolates on *Brugia* adult parasite motility and fecundity using filtrates derived from the specific growth conditions in which they had shown activity in microfilariae (**Figure 1D**). Filtrates of *Pseudomonas viciae* TECH7 significantly reduced adult female motility when the bacteria were grown in M9 and reduced adult male motility when the bacteria were grown in any of the four media. Filtrates from *Pseudomonas viciae* TECH7 (M9 media) and *Pseudomonas* sp. TE4607 (TSB10 media), but not *Pseudomonas* sp. TE4607 (PDB media) reduced adult female fecundity. The distinct phenotypic profiles across bacterial growth conditions suggests that media type significantly influences metabolite production profiles^30^, supporting the relevance of frameworks like One Strain Many Compounds (OSMAC)^31,32^ in nematode screening.

*Pseudomonas* sp. TE4607 was prioritized for follow-up because of its potent effects against *Brugia* microfilariae stage parasites. TSB10 was chosen for culturing because it supported higher activity in the fecundity assay than other media. Effects on *C. elegans* development might be detected with preparations of higher purity, but the results presented here highlight the importance of considering several phenotypic assays in primary screens.

### Labradorin 1 (1) is the primary active compound in *Pseudomonas.* sp. TE4607

Crude extracts from two liters of *Pseudomonas* sp. TE4607 (TSB10) culture were generated by methanol extraction and then partitioned into four solvent phases: chloroform, hexane, n-butanol, and aqueous. These partitions were concentrated to dryness and resuspended in DMSO for screening in the same *C. elegans* and *Brugia* microfilariae assays at a range of concentrations (20 µg/ml - 1 mg/mL, **Figure 2A**). There were low-level activities in all partitions except for the aqueous phase in all three assays (development, motility, and tissue toxicity), but the most potent phenotypes appeared in the hexane partition. HPLC-generated fractions of the three active partitions were screened in the same assays at a final concentration of 100 µg/ml (**Figure 2B**). In these screens, two fractions of the hexane partition were the most active, causing modest reduction in *C. elegans* development and severe reduction in microfilariae motility accompanied by tissue death. These effects were not observed when fractions were screened at 10 µg/ml (data not shown).

**Figure 2.**
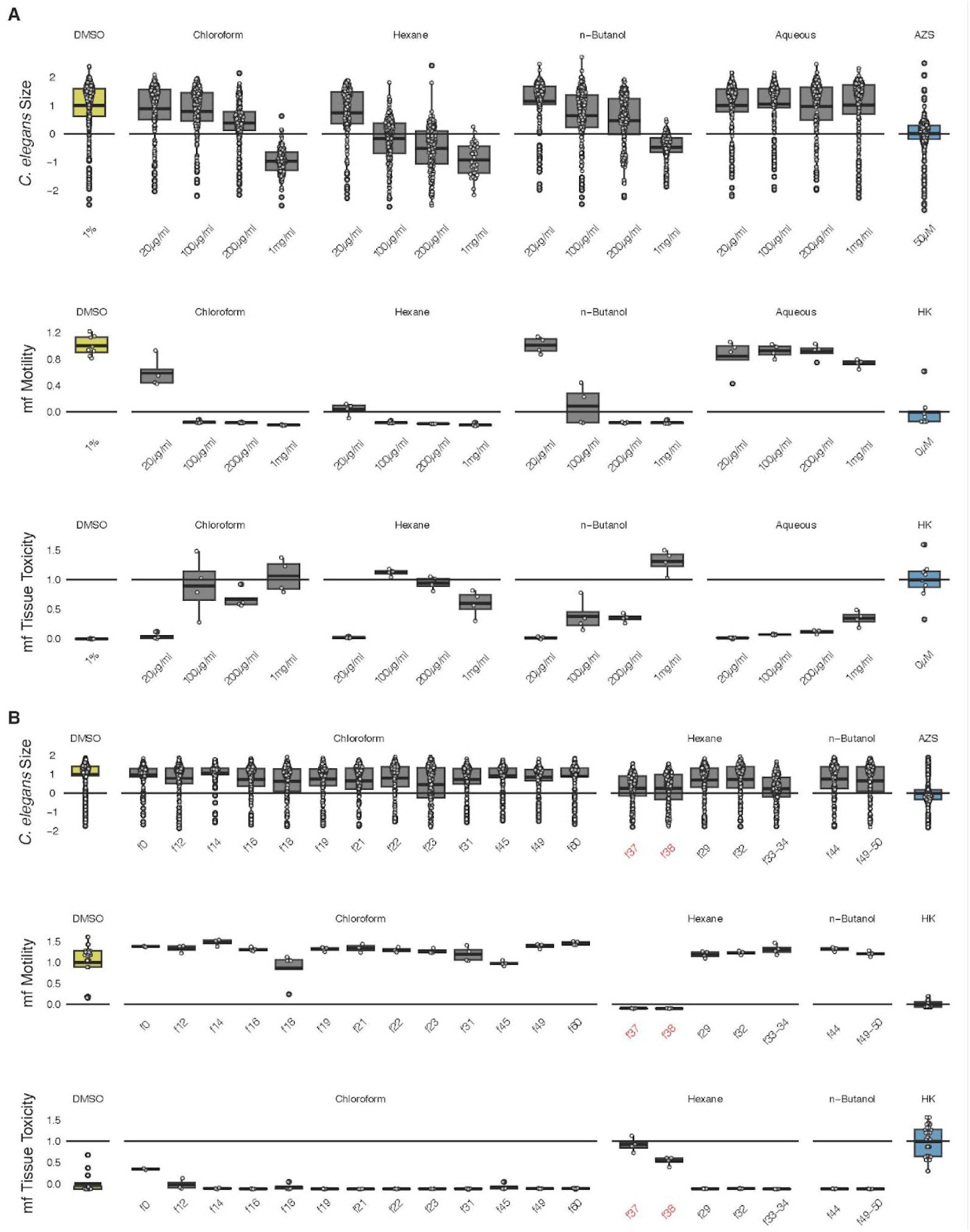
The effects of isolate *Pseudomonas* sp. TE4607 partitions and fractions across nematode phenotypes of interest. **(A)** *C. elegans* development and *Brugia* microfilariae (mf) motility and tissue toxicity endpoints per worm or well after treatment with *Pseudomonas* sp. TE4607 solvent partitions. All values are normalized between mean negative and positive control values. **(B)** *C. elegans* and *Brugia* microfilariae (mf) phenotypes showing individual worm size and total well values of motility and tissue toxicity in the presence of 100µg/ml HPLC fractions generated from each partition. Fractions prioritized for follow-up due to their activity across phenotypes are highlighted in red. Phenotypes were normalized as follows: (X − positive control) / (negative control − positive control) where X is the phenotypic endpoint value.

The chemical structures of active fractions were elucidated using ^1^H, ^13^C NMR, and LCMS (see SI). Compound 1 (Labradorin 1, **Figure 3**) was the only detectable compound in the most active fraction. The second most active fraction contained a mixture of labradorin 1 and pimprinaphine (**Figure 3**). We infer that the lower activity was due to lower abundance of labradorin 1; fractions containing only pimprinaphine were not active at the concentrations tested. Labradorin 1 has previously been isolated from several *Pseudomonas* species^33–35^, and the biosynthetic pathway for synthesis of indolyloxazole alkaloids in this genus has similarly been established^36^. LCMS analysis revealed the previously proposed biosynthetic intermediates (see SI). Labradorin 1 has been reported to be active against certain cancer cell lines^33^ and some species of bacteria^34,35^, but this is the first report of its nematicidal activity. Several derivatives and analogs in the pimprinine family were previously reported to be nematicidal^37^.

**Figure 3.**
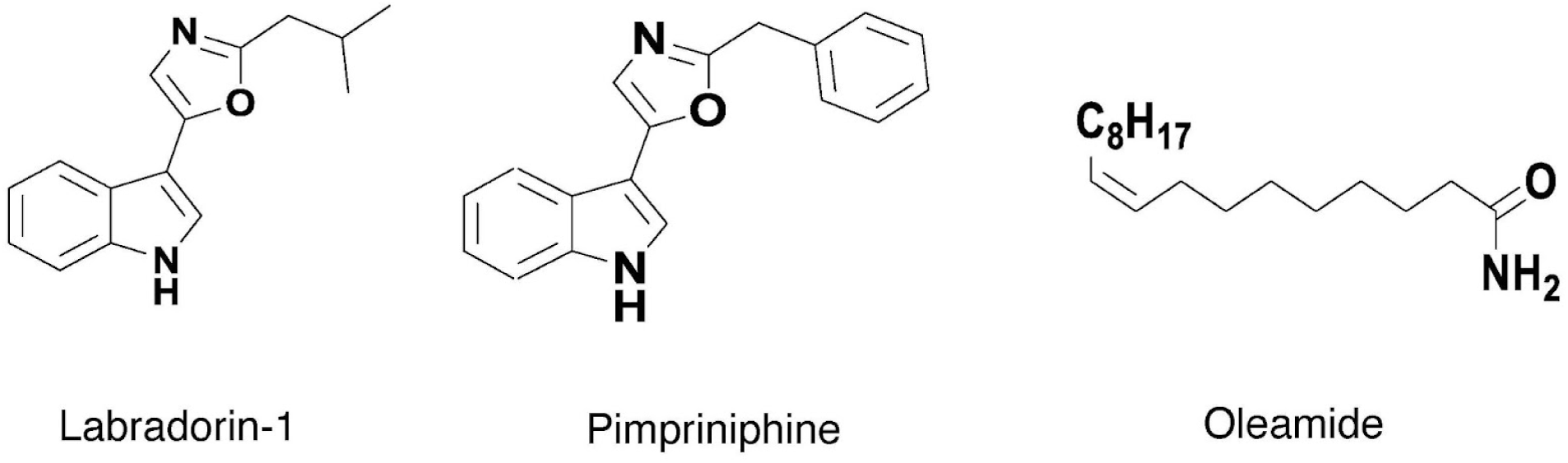
Relevant chemical structures. Labradorin 1(1) was identified as the primary nematicidal active agent from *Pseudomonas* sp. TE4607. Pimpriniphine (2) and oleamide (3) were also identified in active fractions from *Pseudomonas* sp. TE4607 and *Pseudomonas viciae* TECH7.

Extracts from *Pseudomonas viciae* TECH7 grown in M9 had the same profile of activities as TECH4706 grown in TSB10 (see SI). LCMS data indicated the most active compound isolated from the hexane partition was consistent with oleamide (3) (**Figure 3**), alongside trace components of pimprinaphine (2). LCMS data identified labradorin 1 (1) within a mixed-compound fraction that did not exhibit activity, suggesting that other co-occurring compounds may contribute to effects of strain TECH7 on nematodes.

Labradorin 1 (1) was purified from *Pseudomonas* sp. TE4607 culture, and dose-response curves were generated for *Brugia* microfilariae motility (**Figure 4A**) and *C. elegans* development (**Figure 4B**). This resulted in EC50 values of 12.6 µg/ml (52.4 µM) for microfilariae motility and 17.4 µg/ml (72.4 µM) for *C. elegans* development. Labradorin 1 inhibited adult motility at concentrations of 50 µg/ml and above (**Figure 4C**), but did not have a significant effect on adult female fecundity (**Figure 4D**). To determine whether labradorin 1 was toxic to mammalian cells, we tested it on cell line HEK293T, and it was toxic at concentrations near the microfilariae EC50 value (**Figure 4E**). Mammalian toxicity at this level would limit laboradorin’s utility as a monotherapy to treat parasitic nematodes without either chemical modification or a delivery system that reduces host toxicity.

**Figure 4.**
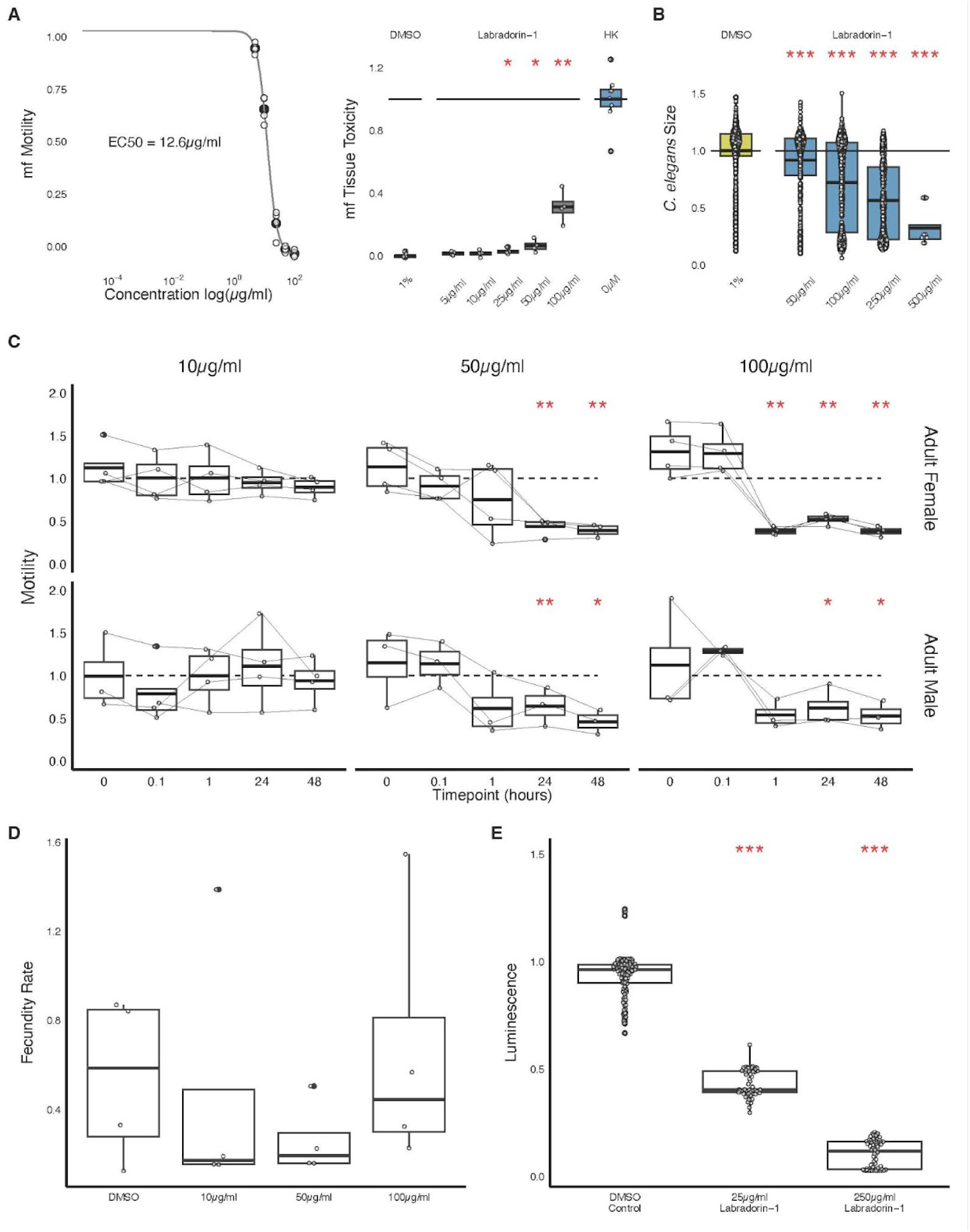
The effects of labradorin 1 on nematodes and mammalian cells. **(A)** *Brugia* microfilariae (mf) motility dose-response curve (left, EC50 = 12.6 µg/ml) and tissue toxicity (right). Values are normalized between mean negative and positive control values. **(B)** *C. elegans* development responses across concentrations (EC50 = 17.3 µg/ml). Values are normalized to mean negative control values. **(C)** *Brugia* adult motility across time points at three labradorin 1 concentrations. Values are normalized to mean negative control values. **(D)** *Brugia* adult fecundity (mf production) at 48 hours post-treatment with three different concentrations of labradorin 1. Values are normalized to initial time point progeny quantities. **(E)** Cell toxicity in Human Embryonic Kidney (HEK293T) cells treated with two concentrations of labradorin-1. A decrease in luminescence indicates cell death and values are normalized to DMSO control values. Statistical analyses were performed via t-test and reported as follows, * : p<0.05, ** : p<0.01, *** : p<0.001, **** : p<0.0001. *C. elegans* and microfilariae phenotypes were normalized as follows: (X − positive control) / (negative control − positive control) where X is the phenotypic endpoint value while adult parasite data points and HEK cell phenotypes were normalized to DMSO and media controls alone: X / negative control.

### Nematode chemoattraction to labradorin 1-producing *Pseudomonas* sp. TE4607

Given the deleterious effects of labradorin 1 on free-living nematodes, we were interested in potential behavioral interactions between these worms and the bacterial strains that produce the compound. Soil nematodes can exhibit chemosensory behaviors in response to metabolites produced by both beneficial and deleterious microbes^38^. For example, the nematode pathogen *Pseudomonas aeruginosa* produces chemoattractants that enable it to infect nematodes^10^. To investigate whether *Pseudomonas* sp. TE4607 modulates sensory responses in *C. elegans*, we used chemotaxis assays to measure nematode attraction to TE4607 supernatant filtrate and purified labradorin 1. In this choice assay (**Figure 5A**), worms were placed in the center of agar plates flanked by test cues and water controls, and the number of worms in each zone was determined over time (1, 2, and 24 hr). This assay was conducted using both an agar-plug soaking method^39^ and a direct application method^40^ to establish chemical gradients.

**Figure 5.**
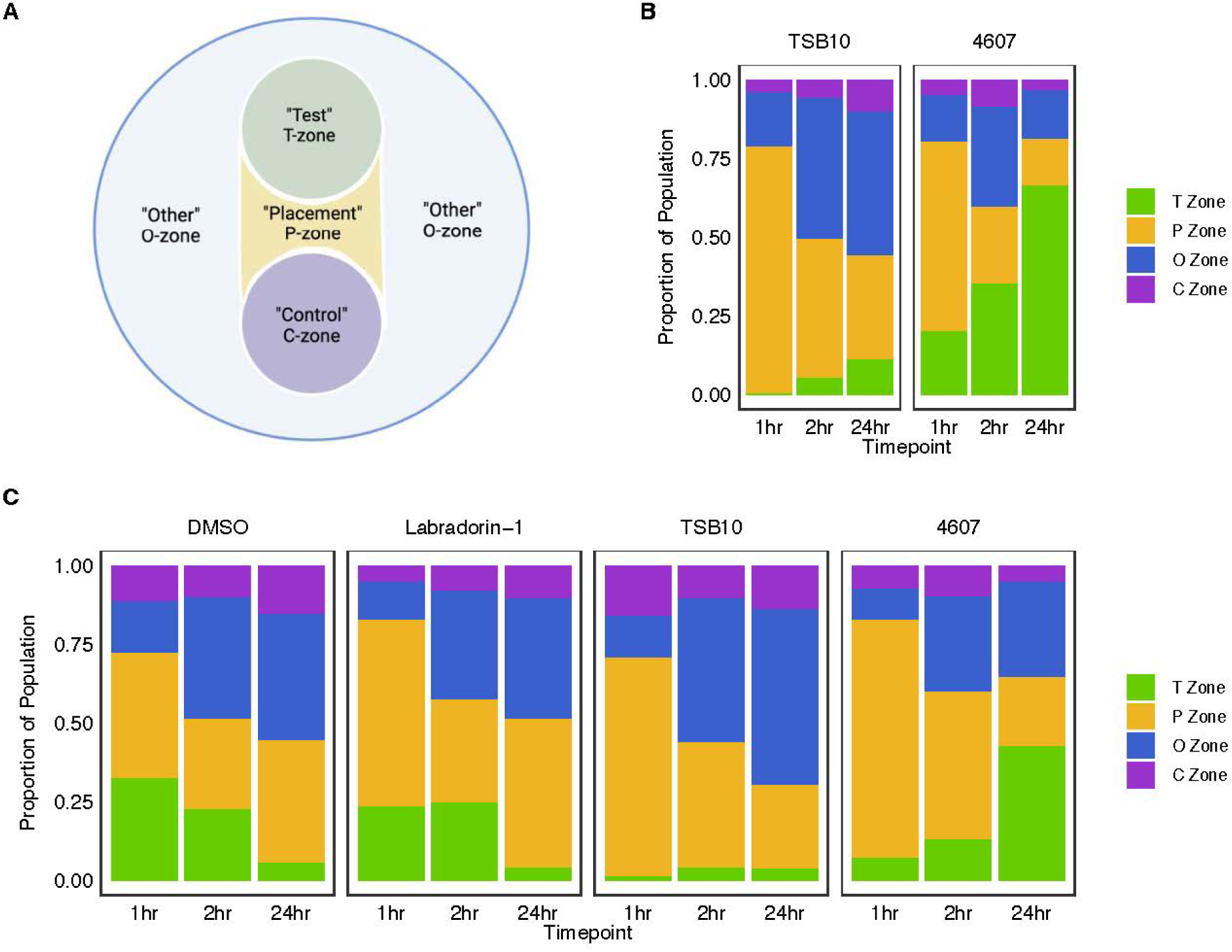
*C. elegans* chemosensory responses to *Pseudomonas* sp. TE4607 filtrate and labradorin 1. **(A)** Depiction of agar plate-based chemosensory choice assay. Worms are placed in the center of the “Placement” zone and treatments are placed by pipette or as soaked agar plugs in the center of the “Test” zone. **(B)** Proportion of worms that are present in each zone over three time points after the T-zone was acclimated with agar plugs soaked in TSB10 control (left) or *Pseudomonas* sp. TE4607 filtrate (right). **(C)** Proportion of worms present in each zone across three time points when the control (DMSO or TSB10) or test cue (Labradorin-1 or *Pseudomonas* sp. TE4607 filtrate) was pipetted onto the center of the T-zone.

Using the agar plug method, we observed strong *C. elegans* attraction towards *Pseudomonas* sp. TE4607 filtrate with most worms accumulating at this cue over the 24-hour observation period. This sensory response is not driven by TSB10 media alone (**Figure 5B**). Next, we used the direct application method, which requires smaller quantities of test cues, to determine whether nematode chemoattraction to *Pseudomonas* sp. TE4607 is decoupled from the filtrate nematicidal activity. The nematodes accumulated around the TE4607 filtrate but populations declined around labradorin 1 over 24 hr, indicating that chemoattraction to TE4607 is independent of labradorin 1 (**Figure 5C**). The transient attraction of worms to labradorin 1 at the earliest time points is explained by the known attraction of *C. elegans* to the solvent, DMSO^41^ in which labradorin was dissolved. Pyoverdin-like compounds might be responsible for chemoattraction because the TE4607 genome contains biosynthetic gene clusters that resemble those of the pyoverdins, which are a known class of *C. elegans* chemoattractants^42^. *Pseudomonas* sp. TE4607 and nematodes have complex chemical interactions, which make the vast pool of soil *Pseudomonas* species a continuously attractive resource for nematicidal compounds for human therapeutics and potential biocontrol agents for plant-parasitic nematodes.

## Experimental Section

### Nematode sources and husbandry

*C. elegans* N2 (Bristol) was maintained on NGM plates seeded with *E. coli* OP50 at 20°C. *Brugia* microfilariae and adult parasites were obtained through the NIH/NIAID Filariasis Research Reagent Resource Center (FR3); morphological voucher specimens are stored at the Harold W. Manter Museum at the University of Nebraska, accession numbers P2021-2032^43^. *Brugia pahangi* and *Brugia malayi* species were used interchangeably according to availability at the time of screening and maintained in RPMI 1640 culture media with penicillin/streptomycin (0.1 mg/ml) at 37°C with 5% atmospheric CO_2_.

### Media and solvents used in this study

Reagent- HPLC-, or LCMS-grade methanol, hexanes, chloroform, *n*-butanol, and acetonitrile were purchased from Fisher Scientific and used as received. Formic acid, trifluoroacetic acid, d4-methanol with TMS internal standard, and HP-20 resin were purchased from Sigma Aldrich and used as received. Tryptic Soy Broth was prepared at 1/10th of the manufacturer’s recommended concentration (TSB10), and Luria-Bertani (LB) and Potato Dextrose Broth (PDB) (BD Bacto™) were prepared according to manufacturer instructions. M9 medium was prepared as follows. A salt stock solution was prepared from anhydrous Na_2_HPO_4_ (33.9 g), KH_2_PO_4_ (15 g), NaCl (2.5 g), and NH_4_Cl (5 g) in 1 L MilliQ water. Salt stock solution (200 mL) was added to 700 mL MilliQ water and autoclaved. Once the solution had cooled to room temperature, sterile 1 M MgSO_4_ (1 mL), 20% m/v glucose (20 mL), and 1 M CaCl_2_ (100 mL) were added per liter of culture broth and the final volume adjusted to 1 L.

### Isolate collection and filtrate preparation

A single colony from solid media of each of the 22 bacterial strains was used to inoculate four different media: LB, TSB10, M9 and PDB. Cultures were shaken at 28℃ at 200 rpm for up to four days. When cultures were turbid (or at 4 days if not turbid), 1 mL was removed from each and stored at −20℃. Cells and debris were removed from the remaining culture by centrifugation, and the resulting supernatant was filtered through a 0.2-μm filter and frozen until use.

Isolate species was first determined by the IMG annotation pipeline^44^ (NCBI tax id: d_Bacteria; p_Pseudomonadota; c_Gammaproteobacteria; o_Pseudomonadales; f_Pseudomonadaceae; g_Pseudomonas; s_Pseudomonas hunanensis) and later classified by GTDB-tk^45^ (d_Bacteria;p_Pseudomonadota; c_Gammaproteobacteria; o_Pseudomonadales; f_Pseudomonadaceae;g_Pseudomonas_E; s_Pseudomonas_E sp024749165). *Pseudomonas* sp. TE4607 was isolated by students at Beloit College from a soil sample taken in Beloit, Wisconsin. *Pseudomonas viciae* TECH7 was isolated by students at the University of Northwestern, St. Paul from a soil sample taken in Maplewood, Minnesota.

### Extract, partition, and fraction preparation

#### Pseudomonas sp. TE4607 culture and chemical isolation conditions

A three-way streak plate was prepared on LBA from a glycerol stock stored at −80°C and incubated at 28°C for two days. Sterile LB broth (5 mL) was inoculated with a single colony and incubated overnight at 28°C. TSB10 (2×1 L) was inoculated with overnight culture (1 mL/L) and the broth incubated at 28°C in a shaking incubator at 200 RPM for 24 hours. Pre-activated HP-20 resin (70 g/L) was added and the broth culture shaken for an additional hour. The resin was collected by filtration through miracloth and washed with water (3×500 mL), transferred to a large beaker, and extracted with methanol for one hour in triplicate (3×300 mL). The combined extracts were concentrated under reduced pressure to gain the crude extract. The crude extract was resuspended in 10% MeOH/H_2_O (200 mL) with sonication, and sequentially partitioned into hexanes (3×50 mL), chloroform (3×50 mL), and *n*-butanol (3×50 mL). The initial partitioning into hexanes often produced an emulsion, which was dispersed using a minimal amount of brine. Each organic layer was washed with water (2×20 mL) and brine (20 mL), dried over Na_2_SO_4_, filtered, and concentrated under reduced pressure. The partitioned extracts were resuspended in methanol, filtered through a 0.2 mm PTFE filter, and the solvent removed. The hexanes and chloroform partitions were subjected to chromatographic separation by reverse phase HPLC. HPLC analyses were performed on a Shimadzu Nexera Series with a PDA and ELSD detector. A Phenomenex™ Luna 5 mm C18 column with dimensions of either 4.6×250 mm or 10×250 mm was used for analytical and semi-prep scale separations, respectively. HPLC chromatograms were processed using LabSolutions software. ^1^H and ^13^C NMR spectra were measured on a Bruker Avance-500 equipped with a DCH cryoprobe or a Bruker Avance-400 equipped with a BBFO probe. NMR spectra were processed using MestraNova software. All chemical shifts are reported in units of parts per million (ppm) downfield from tetramethylsilane (TMS) and coupling constants are reported in units of hertz (Hz). High resolution mass spectra were measured on a Thermo Q Exactive Plus™ using electrospray ionization in tandem with a Vanquish VH-P10 LC system equipped with a Phenomenex™ kinetex 1.7 mm C18 column with dimensions of 2.1×100 mm. LCMS spectra were processed using FreeStyle software. Compound Discoverer (version 2.0, Thermo Fisher) was used for assigning known compounds with documented mass spectra and cross checked using Natural Products Atlas and SciFinder where possible. Purification by HPLC (10-90% or 10-30-70% MeCN/H_2_O with 0.1% TFA over 30 min; 4.7 ml/min) provided several purified compounds and characterization data are described in supplemental materials.

*Pseudomonas viciae* TECH7 was cultured similarly to *Pseudomonas* sp. TE4607 in M9 broth and TSB10, respectively. Chemical extraction and purification followed the same procedure as described above.

### Nematode screening protocols

Isolate and extract samples were stored at −20°C and thawed, diluted, and aliquoted to empty 96-well assay plates (Greiner Bio-One 655180) immediately prior to assay setup. Filtrate samples were added to plates in volumes of 10 µl per well (1:10 dilution). Extract samples were stored dry and diluted using DMSO to 100X tested concentrations based on extract or fraction weight. Resuspended samples were added to plates in volumes of 1µl per well (1:100 dilution). Media alone was used as a negative control in place of DMSO for filtrates. Nematodes were prepared according to species and assay as follows. ***C. elegans development:*** *a*pproximately 18 hours prior to development screening, gravid worms were synchronized via bleaching^46^, and embryos were hatched in filter-sterilized K media^47^. Titering of larvae, preparation of food mixture, and set up and incubation of 96-well assay plates were performed as previously described^23^. After 48 hours, assay plates were rinsed with M9 using an AquaMax 2000 plate washer (Molecular Devices), and sodium azide (Thermo Scientific) was added at a final concentration of 50mM to paralyze worms. Whole wells were imaged with a 2X objective using an ImageXpress Nano (Molecular Devices), and images were analyzed using the worm size module of wrmXpress v1.4.0^27^. ***Brugia microfilariae*:** motility and tissue toxicity assays using CellTox Green (Promega) were performed as previously described^48^. Images were acquired using a 4X objective on an ImageXpress Nano and analyzed using the motility and cell toxicity modules of wrmXpress^27^. ***Brugia adult parasite:*** motility and fecundity assays were set up as described^49^ with minor modifications. Parasites were transferred between plates after the 0-hour and 48-hour time points. Motility videos were cropped using Fiji^50^ and analyzed using optical flow^23^. Fecundity images were stitched and segmented using a previously developed Fiji protocol^49^. All endpoints were normalized and analyzed using R software including tidyverse packages^51^ for statistical analysis and the drc package^52^ for dose-response analyses. *C. elegans* and microfilariae phenotypes were normalized as follows: (X − positive control) / (negative control − positive control) where X is the phenotypic endpoint while adult parasite data points were normalized to DMSO and media controls alone: X / negative control after normalizing to individual initial motility scores.

### Cell Line Toxicity Screening

HEK293T cells were cultured in DMEM high glucose + GLUTAMAX + pyruvate (Life technologies, 10569010) supplemented with 10% FBS (Fisher A52567) and penicillin-streptomycin (Cytiva SV30010) at 100 U/mL and 100 µg/mL, respectively. For maintenance, cells were split when 80% confluence was reached and passed at a split ratio between 1:5 and 1:10. Briefly, cells were washed with DPBS without calcium or magnesium (Gibco, 14190144), trypsinized with 0.05% trypsin-EDTA (Gibco, 25300054), resuspended in culture media, and passed to a T25 flask with fresh culture media. For toxicity assay set up, cells were plated in culture media with dialyzed FBS (Cytiva, SH30079.02) in white, opaque plates (Greiner Bio-One, 07-000-138) 48 hours in advance of drug application. Cells were plated at a concentration that yielded 80% confluence the day of the assay. Some wells contained media with no cells as controls and to cell wells, DMSO or labradorin-1 was added at a 1:100 dilution. After 24 hours, plates and CellTiter-Glo Luminescent Cell Viability Assay (Promega) reagent were equilibrated to room temperature before the reagent was added to wells according to label instructions. Plates were then left at room temperature and protected from light exposure for 10 minutes prior to using a SpectraMax Plate Reader (Molecular Devices) to read luminescence values using an integration time of 750ms. Luminescence data was analyzed using R software and normalized to DMSO controls.

### C. elegans chemosensory assay

Chemotaxis agar media^53^ was prepared (2.5% agar, 1mM CaCl_2_, 5mM KHPO_4_, 1mM MgSO_4_) and poured into 10 cm petri dishes. Plate markings were drawn based on previous chemotaxis screens performed for filarial nematodes^54,55^ with slight modifications to accommodate the plate size. Briefly, two circles (25mm diameter) sat on opposing sides of the plate, each sitting 0.5 cm from the plate edge, and the midpoint of the plate was marked. *Pseudomonas* sp. TE4607 supernatant was produced by cultivating *Pseudomonas* sp. TE4607 in TSB10 media and then manually separating cells from the supernatant through repeated centrifugation. Two different approaches were adapted to create cue gradients on the assay plates based on *C. elegans* chemotaxis studies using agar plug^39^ and direct spotting^40^ methods. First, plugs were cut from a plate using the large end of a 1000 µL pipette tip and soaked for 5 hours in a microcentrifuge tube filled with 1 ml of media or supernatant and rotated on a nutator. Soaked plugs were then set in the center of one plate circle (T-zone) overnight, and removed from plates immediately before adding worms. For direct spotting, 5 µl of test sample (*Pseudomonas* sp. TE4607 supernatant, TSB10, DMSO, or 10 mg/ml labradorin-1) were added to the middle of the T-zone circle while 5 µl of MilliQ water (negative control) was added to the middle of the C-zone circle and allowed 3 hours to disperse. *C. elegans* were prepared by picking 5 L4 worms to several 6 cm plates and incubating for 4 days. On the day of assay set up, *C. elegans* were collected from maintenance plates with M9 media and washed twice with M9 and once with MilliQ water before being resuspended in 1 mL of MilliQ water and counted. A volume equivalent to ~150 adult worms was added to the center point of the plate. The worms in each of the zones were counted manually at 1, 2, and 24 hours after *C. elegans* were transferred to plates and able to chemotax. For agar plug assays, one plate was used per drug condition while for direct spotting assays, two plates per drug condition were performed.

## Acknowledgments

Support for this research was provided by the University of Wisconsin-Madison Office of the Vice Chancellor for Research and Graduate Education with funding from the Wisconsin Alumni Research Foundation. Additional support was provided by National Institutes of Health NIAID grant R01 AI151171 to MZ and the Department of Education grant P116Z230024 to JH. KTR was funded by the National Institutes of Health grant 5T32AI007414. *Brugia* life cycle stages were obtained through the NIH/NIAID Filarial Research Reagent Resource Center (FR3), morphological voucher specimens are stored at the Harold W Manter Museum at University of Nebraska, accession numbers P2021-2032. Finally, the authors would like Elena Rehborg for maintaining and preparing the HEK293T cell line used for mammalian cell toxicity screens. Support for MGC was provided by the Department of Plant Pathology at the University of Wisconsin-Madison and a Research Starter Grant from the American Society for Pharmacognosy.

## Notes

### Competing Interest Statement

The authors have declared no competing interest.

