## Supplemental Materials for "Nematicidal indole oxazoles and chemoattractants from soil bacteria"

### **Compound characterization**

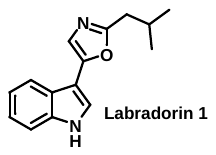

Labradorin 1 (numberX). Isolated as a white solid (1.7 mg) from the hexanes partition obtained from 2L *P. soli* TE4607 culture. The structure of labradorin 1 was confirmed by ^1^H, ^13^C NMR, and LCMS. Spectral data agreed with reported values. ^refs^ ^1^H NMR (500 MHz, CDCl_3_) δ 7.82 (dd, *J* = 7.9, 1.1 Hz, 1H), 7.64 (s, 1H), 7.47 (dd, *J* = 7.9, 1.1 Hz, 1H), 7.27 – 7.21 (m, 2H), 7.18 (ddd, *J* = 8.0, 7.0, 1.2 Hz, 1H), 2.78 (d, *J* = 7.2 Hz, 2H), 2.31 – 2.18 (m, 1H), 1.07 (d, *J* = 6.7 Hz, 5H). ^13^C NMR (126 MHz, CDCl_3_) δ 163.72, 150.26, 138.19, 125.32, 123.87, 123.47, 121.39, 120.41, 118.56, 112.84, 105.31, 37.68, 28.93, 22.62. HRMS (ESI) m/z (%): cald. for C_15_H_17_N_2_O [M+H] 241.1340; found 241.1333; 241.1333 (100%), 130.0651 (29.5%), 57.07011 (25.2%), 157.0759 (7.6%), 169.0758 (5.7%), 198.0786 (4.5%), 85.0647 (3.1%).

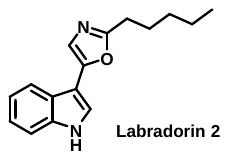

Labradorin 2 (numberX). Isolated as a white solid (0.2 mg) from the hexanes partition obtained from 2L *P. soli* TE4607 culture. The structure of numberX was confirmed by LCMS. Spectral data agreed with reported values. ^refs^ . HRMS (ESI) m/z (%): cald. for C_16_H_19_N_2_O [M+H] 255.1497; found 255.1490: 255.1491 (100%), 130.0651 (29.1%), 71.08553 (11.5%), 157.0759 (8.8%), 103.0542 (2.7%), 169.0758 (2.2%), 99.08038 (1.9%), 198.0786 (1.9%).

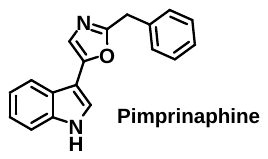

Pimprinaphine (numberx). Isolated as a pale yellow oil (trace) from the hexanes partition obtained from 2L *P. soli* TE4607 culture. The structure of numberX was confirmed by LCMS. Spectral data agreed with reported values. ^refs^ HRMS (ESI) m/z (%): cald. for C_18_H_15_N_2_O [M+H] 275.1184; found 275.1178: 275.1178 (100.0%), 91.0542 (98.4%), 142.0651 (21.4%), 130.0651 (21.0%), 115.0542 (16.9%), 157.0759 (9.1%), 65.0387 (8.7%), 230.0962 (7.6%), 197.0709 (7.4%), 103.0541 (6.2%), 220.1116 (3.1%).

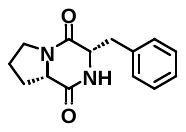

Cyclo(L-phenyl-L-prolyl) (numberx). Isolated as a pale yellow oil (0.5 mg) from the chloroform partition obtained from 2L *P. soli* TE4607 culture. The structure of numberX was confirmed by LCMS. Spectral data agreed with reported values. ^refs^ HRMS (ESI) m/z (%): cald. for C_14_H_17_N_2_O_2_ [M+H] 245.1290; found 245.1284: 70.0652 (100.0%), 245.1284 (79.7%), 120.0807 (70.5%), 98.0600 (29.3%), 154.0735 (18.0%), 103.0542 (15.6%), 217.1333 (6.0%), 91.0542 (4.2%), 172.1119 (3.9%), 153.0657 (3.3%).

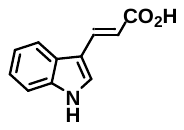

(E)-3-(1H-indol-3-yl)acrylic acid (numberX). Isolated as a white solid (0.4 mg) from the *n*-butanol partition obtained from 2L *P. soli* TE4607 culture. The structure of numberX was confirmed by LCMS. Spectral data agreed with reported values. ^refs^ HRMS (ESI) m/z (%): cald. for C_11_H_10_NO_2_ [M+H] 188.0711; found: 188.0704; 146.0599 (100.0%), 118.0650 (68.4%), 188.0704 (36.9%), 91.0541 (33.8%), 115.0541 (26.7%), 144.0807 (23.1%), 143.0729 (16.8%), 117.0572 (12.3%), 142.0651 (10.6%), 170.0598 (10.6%), 117.0698 (4.4%), 116.0621 (3.1%).

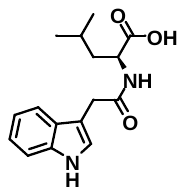

(2-(1H-indol-3-yl)acetyl)-L-leucine (numberX). Isolated as a yellow oil from the *n*-butanol partition obtained from 2L *P. soli* TE4607 culture. The structure of numberX was confirmed by LCMS. Spectral data agreed with reported values. ^refs^ HRMS (ESI) m/z (%): cald. for C_16_H_21_N_2_O_3_ [M+H] 289.1552; found: 289.1543; 188.0704 (100.0%), 205.0970 (74.9%), 146.0599 (56.9%), 159.0915 (43.6%), 118.0649 (37.8%), 57.0701 (20.0%), 115.0540 (18.4%), 130.0650 (17.7%), 132.0807 (16.1%), 91.0541 (14.8%), 243.1489 (14.0%), 117.0571 (13.4%), 144.0806 (10.7%), 143.0728 (9.2%), 85.0647 (7.0%), 142.0652 (6.9%), 170.0598 (6.9%), 289.1543 (5.3%), 187.0866 (3.9%).

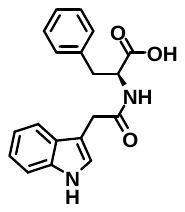

(2-(1H-indol-3-yl)acetyl)-L-phenylalanine (numberX). Isolated as a yellow oil from the *n*-butanol partition obtained from 2L *P. soli* TE4607 culture. The structure of numberX was confirmed by LCMS. Spectral data agreed with reported values. ^refs^ HRMS (ESI) m/z (%): cald. for C_19_H_19_N_2_O_3_ [M+H] 323.1395; found: 323.1382; 91.0541 (100.0%), 188.0704 (80.6%), 159.0915 (60.2%), 146.0599 (47.7%), 277.1334 (45.3%), 205.0969 (40.2%), 130.0650 (35.5%), 118.0650 (32.0%), 115.0541 (23.5%), 132.0807 (23.1%), 117.0571 (16.4%), 170.0597 (10.0%), 144.0806 (8.8%), 142.0649 (8.1%), 143.0728 (7.9%), 65.0387 (6.7%), 187.0864 (5.2%), 305.1281 (4.8%), 323.1382 (4.6%), 160.0757 (3.1%).

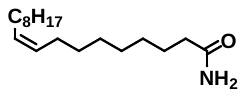

Oleamide (numberX). Isolated as a colorless wax (0.3 mg) from the hexanes partition obtained from 2L *P. viciae* TECH7 culture. The structure of numberX was confirmed by LCMS. Spectral data agreed with reported values. ^refs^ HRMS (ESI) m/z (%): cald. for C_18_H_36_NO [M+H] 282.2796; found: 282.2786; 282.2786 (100.0%), 96.0805 (15.3%), 98.0598 (14.6%), 264.2681 (11.6%), 99.0676 (10.5%), 82.0649 (5.9%), 55.0544 (4.0%), 110.0961 (3.7%)

### **NMR and MS/MS Spectra**

#### Labradorin 1 (numberX) ^1^H NMR (500 MHz, MeOH-d_4_)

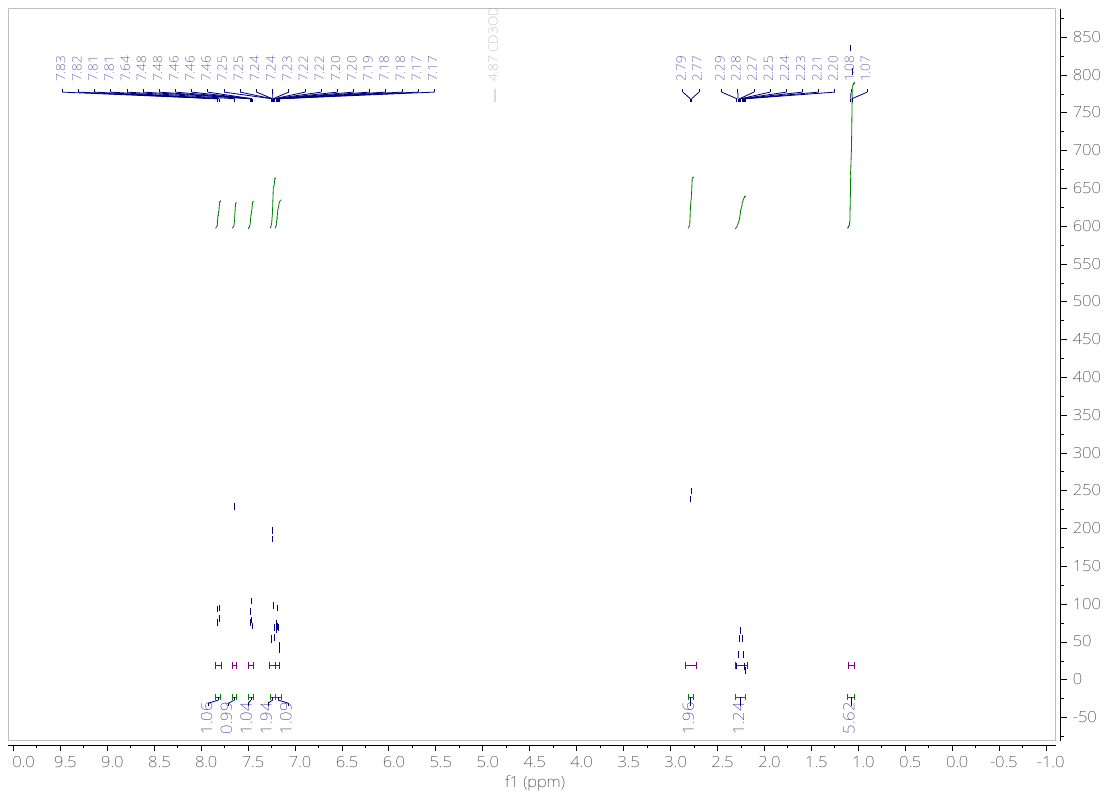

#### Labradorin 1 (numberX) ^13^C NMR (126 MHz, MeOH-d_4_)

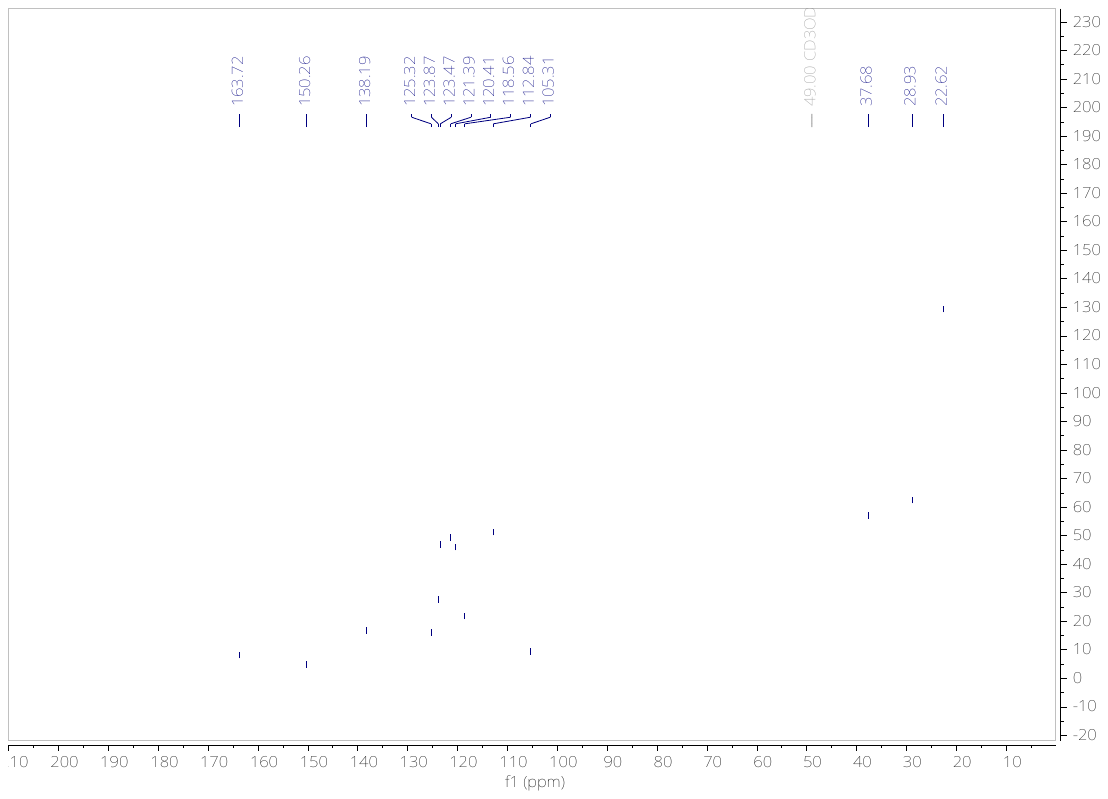

#### Labradorin 1 (numberX) MS/MS, ESI (Orbitrap)

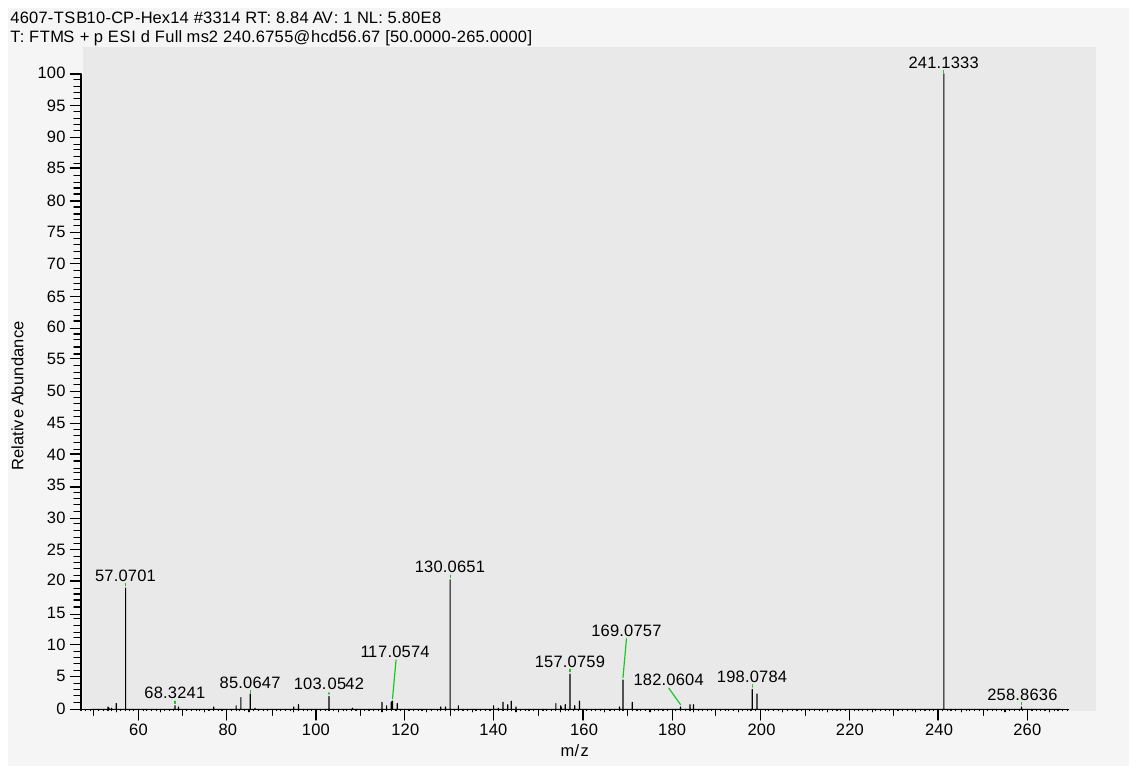

#### Labradorin 2 (numberX) MS/MS, ESI (Orbitrap)

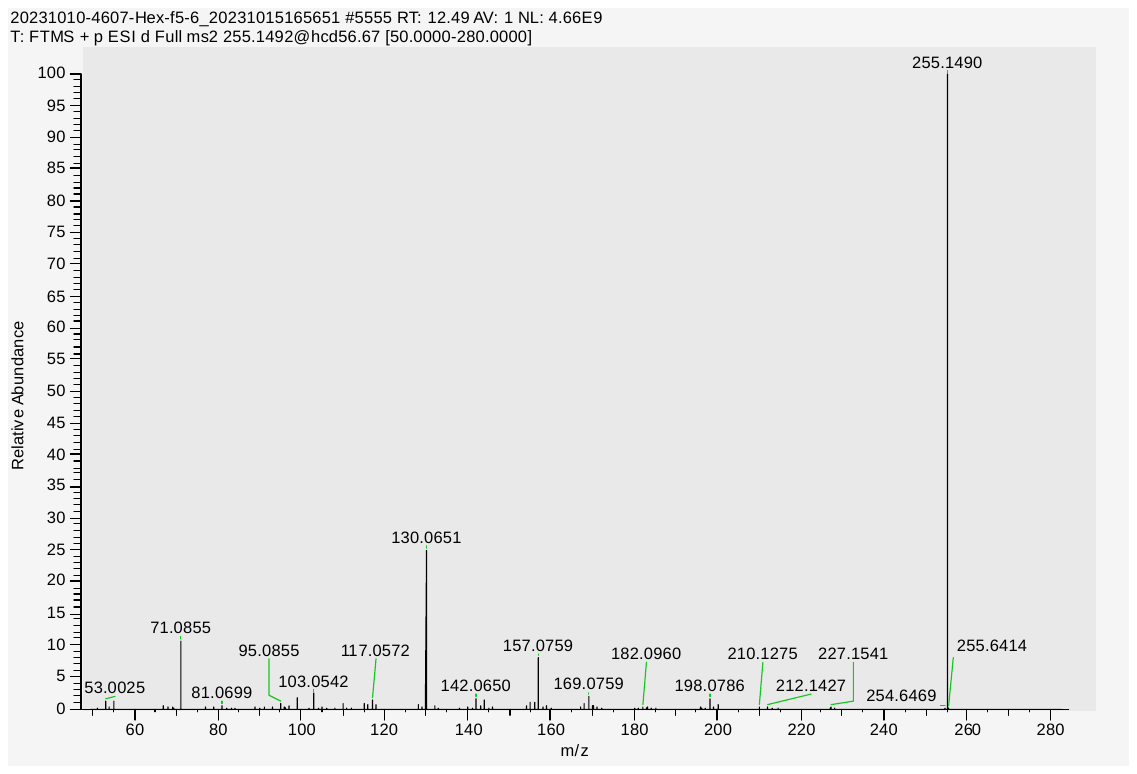

#### Pimprinaphine (numberX) MS/MS, ESI (Orbitrap)

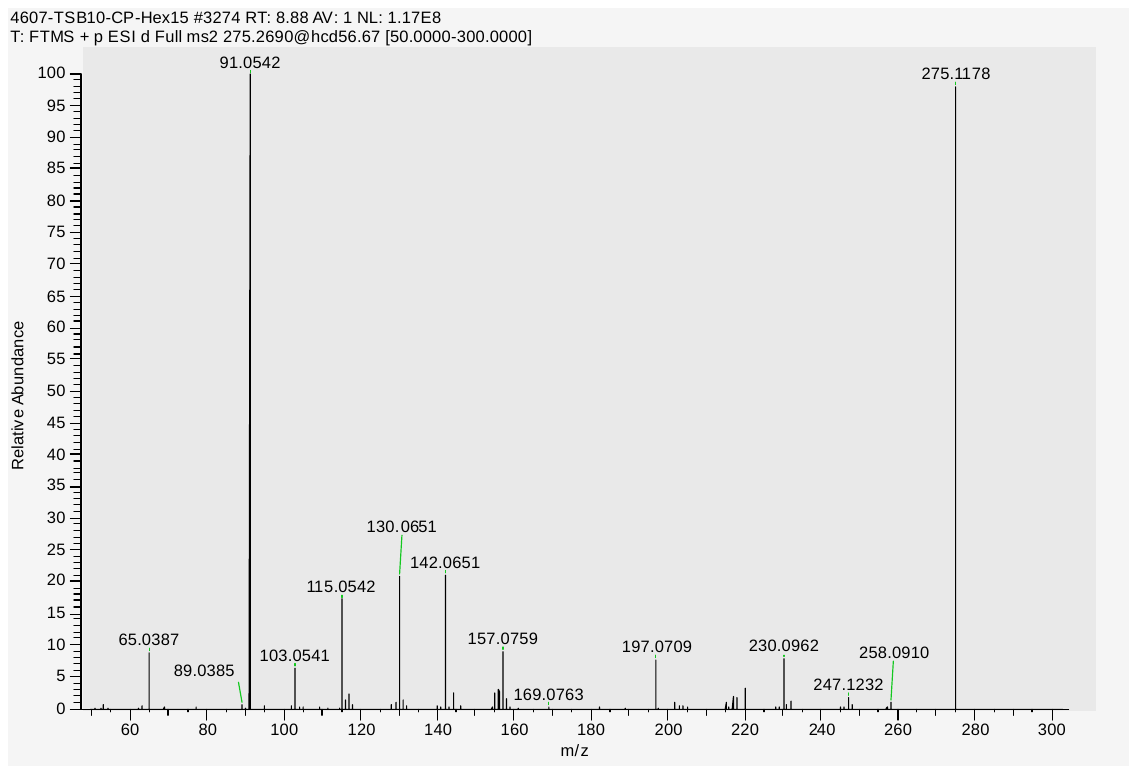

#### Cyclo(L-phenyl-L-prolyl) (numberX) MS/MS, ESI (Orbitrap)

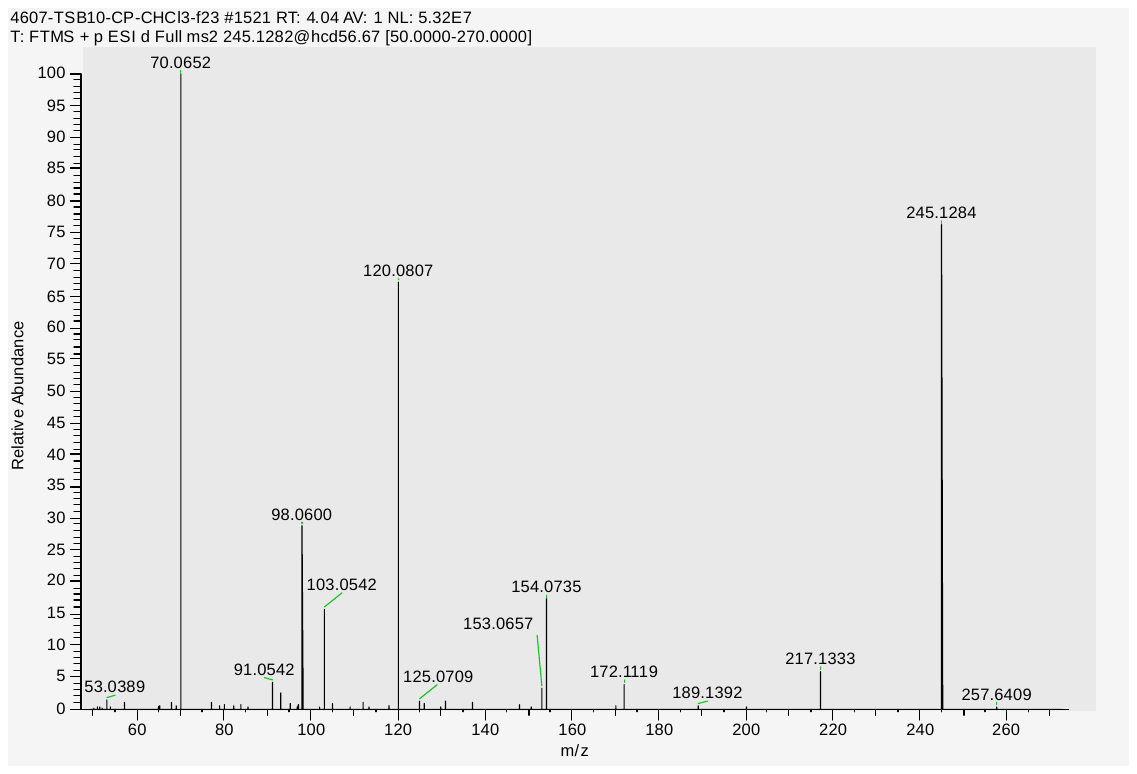

#### (2-(1H-indol-3-yl)acetyl)-L-leucine (numberX) MS/MS, ESI (Orbitrap)

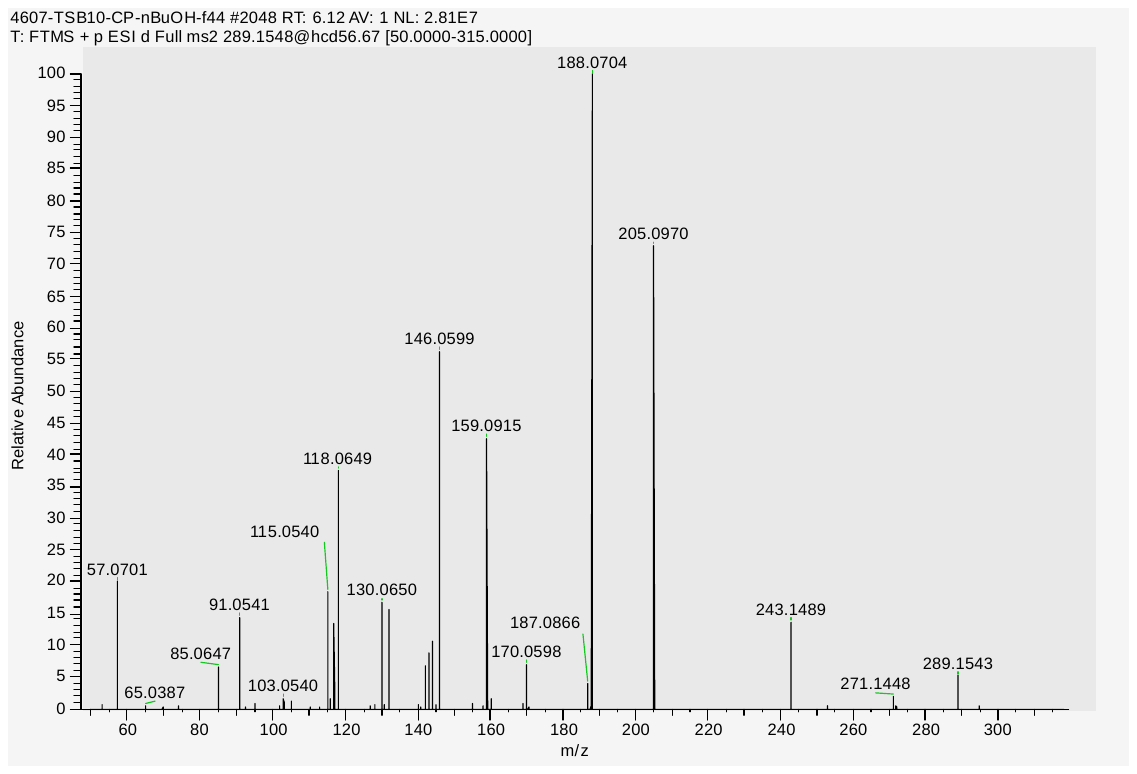

#### (2-(1H-indol-3-yl)acetyl)-L-phenylalanine (numberX) MS/MS, ESI (Orbitrap)

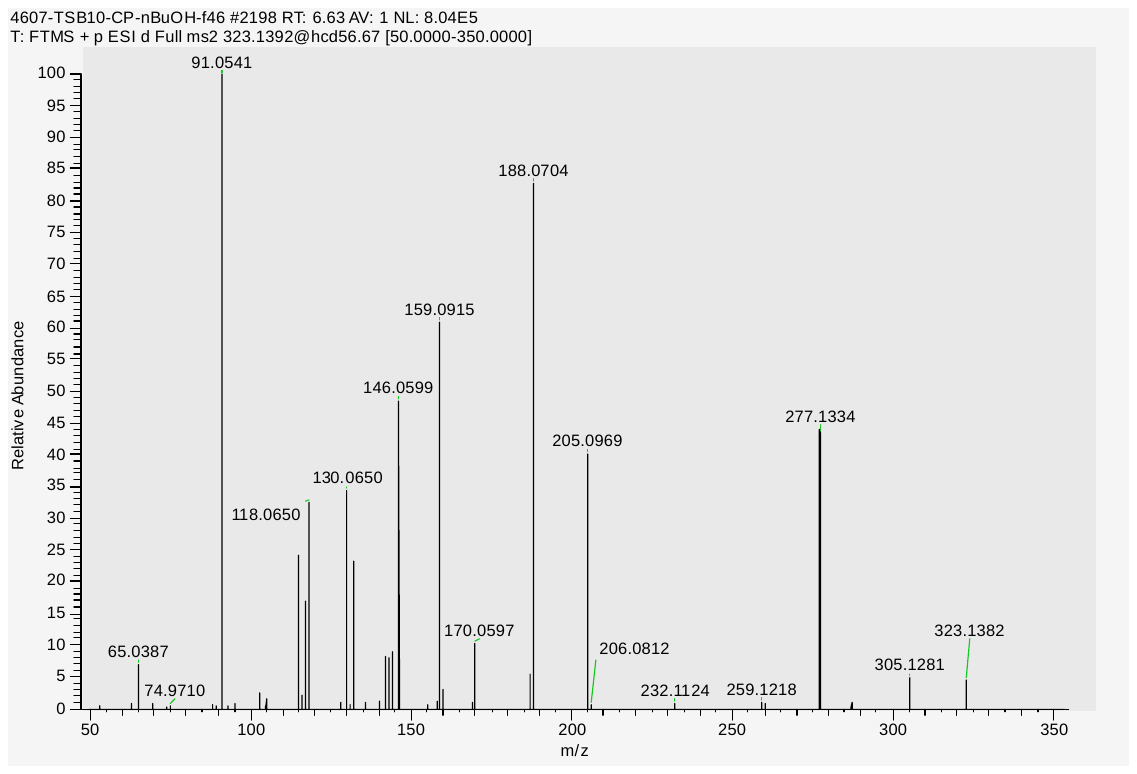

#### (E)-3-(1H-indol-3-yl)acrylic acid (numberX) MS/MS, ESI (Orbitrap)

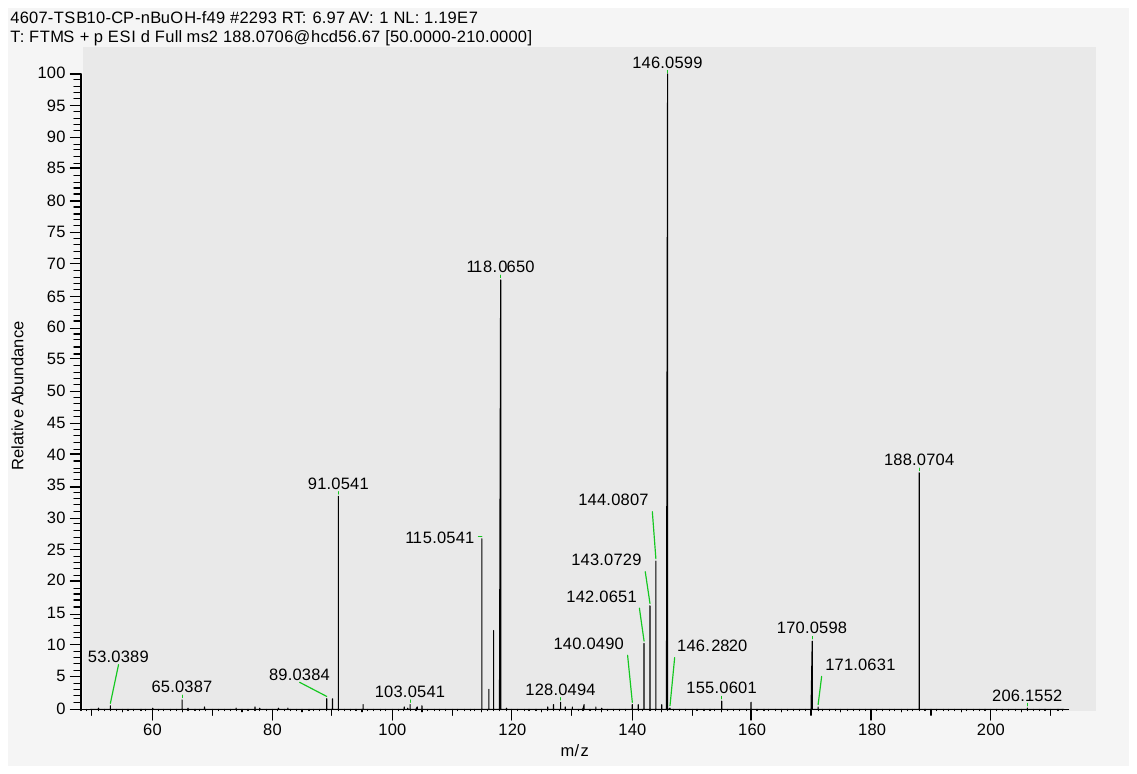

#### Oleamide (numberX) MS/MS, ESI (Orbitrap)

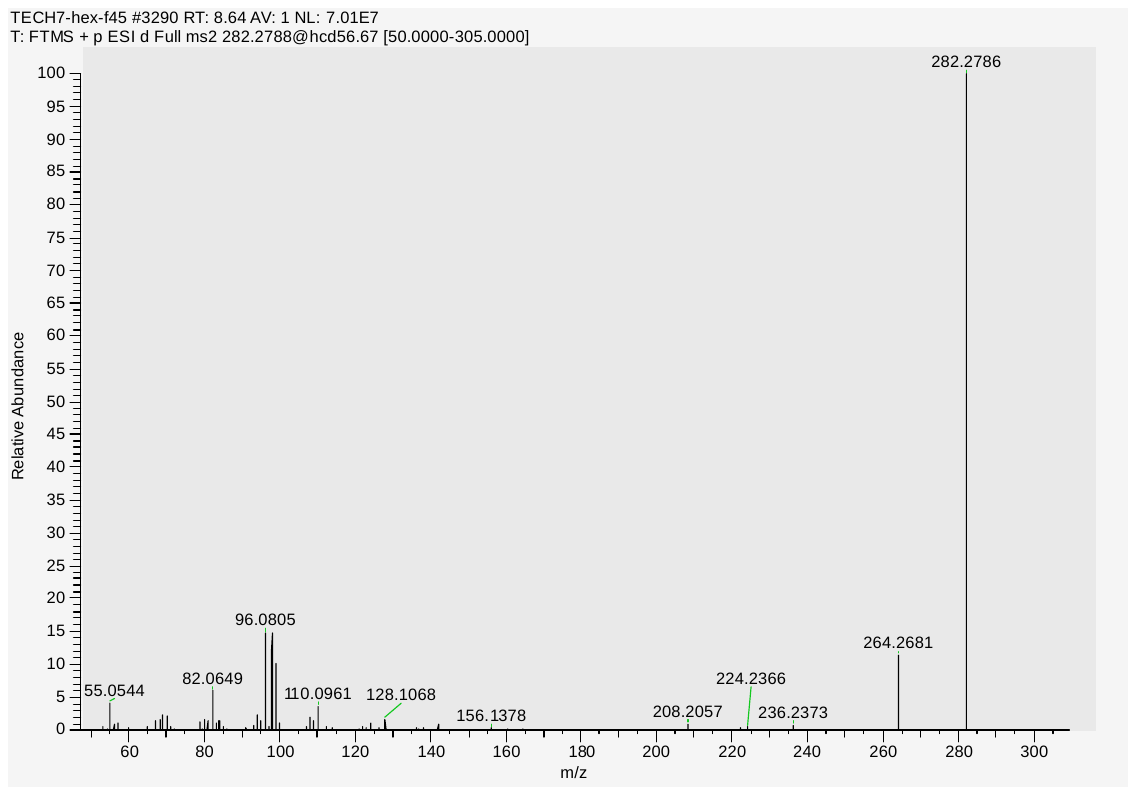
